# MicroRNAs are likely part of the molecular toolkit regulating adult reproductive diapause in the mosquito, *Culex pipiens*

**DOI:** 10.1101/392738

**Authors:** Megan E. Meuti, Robin Bautista, Julie A. Reynolds

## Abstract

For many arthropods, including insects, diapause is the primary mechanism for survival during unfavorable seasons. Although the exogenous signals and endogenous hormones that induce and regulate diapause are well-characterized, we still lack a mechanistic understanding of how environmental information is translated into molecular regulators of the diapause pathway. However, short, non-coding microRNAs (miRNAs) are likely involved in generating both the arrested egg follicle development and fat hypertrophy in diapausing females of the Northern house mosquito, *Culex pipiens*. To determine whether miRNAs might respond to changes in day length and/or regulate diapause pathways, we measured the abundance of candidate miRNAs in diapausing and nondiapausing females of *Cx. pipiens* across the adult lifespan. Of the selected miRNAs nearly all were more abundant in nondiapausing females relative to diapausing females, but at different times. Specifically, miR-13b-3p, miR-14-3p, miR-277-3p, and miR-305-5p were upregulated in nondiapausing females early in adulthood, while miR-309-3p and miR-375-3p were upregulated later in adult life, and miR-8-3p and miR-275-3p were upregulated both early and late in adult life. Taken together, our data demonstrate that miRNA expression is dynamic, changing across adult lifespan. Further, differential miRNA expression between diapausing and nondiapausing females of *Cx. pipiens* suggests that this epigenetic mechanism is part of the molecular toolkit regulating diapause.

## 1. Introduction

Diapause is an alternative developmental pathway that provides insects, and other animals, a means to survive periods of inimical environmental conditions and exploit seasons with abundant resources. Diapause is a dormant state characterized by developmental arrest, metabolic depression, and enhanced tolerance of environmental stresses. This anticipated response is endogenously regulated and induced during a genetically determined stage of the life-cycle in response to token cues (e.g. changes in photoperiod, temperature, or food quality) that signal the advent of unfavorable conditions. Perception of the appropriate cues promotes changes in gene regulatory networks that ultimately lead to physiological changes that define the diapause phenotype [1]. Ecologically relevant token cues that induce the diapause program are known for many insects [2–5], and diapause-relevant changes in gene expression have been documented for a variety of arthropod species [6–13]. Furthermore, many aspects of diapause have been particularly well-characterized in several mosquito species (reviewed in [14],[15]). However, many details of the regulatory networks that mediate the diapause response remain undefined. A particular knowledge gap is how epigenetic processes, defined here as any process that can alter the phenotype independent of changes to the genotype, may regulate diapause induction, maintenance, and/or termination. The goal of the current study was to explore the role of microRNAs (miRNAs), one type of epigenetic mechanism, in regulating reproductive diapause in adult females of the Northern house mosquito, *Culex pipiens*.

MiRNAs are small (18–25 nucleotides) non-coding RNAs that post-transcriptionally regulate gene expression though their interactions with target gene transcripts. Mature miRNA sequences, processed from the 3′ or 5′ arms of a single-stranded hairpin precursor, bind to Argonaut 1 (Ago1) within the RNA Induced Silencing Complex (RISC) and guide the complex to target mRNAs. Once bound to the RISC, miRNAs can increase expression of their targets [16–19]; but more commonly, miRNAs silence target genes via transcript degradation or translation repression. MiRNAs are emerging as regulators of diapause and other dormant states. They have been implicated as regulators of numerous diapause-relevant biological processes including cell-progression [20, 21], developmental timing [22, 23], metabolism [24, 25], and stress-resistance [26–28]. And, a number of recently published studies have identified miRNAs that are differentially regulated in diapausing insects and other animals including diapausing embryos of the crustacean *Artemia parthenogenetica* [21], the killifish *Austrofundulus limnaeus* [29], and insects including the flesh fly *Sarcophaga bullata* [30] and the mosquito *Aedes albopictus* [31]. Taken together, the accumulating evidence on miRNA function and abundance in diapausing animals suggests that miRNAs are part of the “diapause toolkit” [9] that regulates diapause across species.

The goal of the present study is to evaluate changes in the abundance of candidate miRNAs in diapausing adults of a *Culex pipiens*, a mosquito that is an established model for studying adult-reproductive diapause. Female adults of this species survive the winter by entering diapause approximately 5 days after adult emergence in response to short day lengths (< 12 h light per 24 h) received during the 4^th^ larval instar, pupal and early adult stages of development [32, 33]. The diapause phenotype in this species is characterized by a lack of host-seeking behavior, arrested ovarian development, fat hypertrophy (i.e., accumulated lipid stores), suppressed metabolism, and enhanced resistance to stress from low temperatures, desiccation, and pathogens [34–39]. In addition, the roles of juvenile hormone and insulin signaling in establishing and maintaining diapause have been well described for this species [40–41], and components of the circadian clock have been implicated as regulators of diapause in this species [42]. This knowledge about the regulation of diapause, as well as information provided about the biological functions of numerous miRNAs in other mosquito species [31, 43–47] makes *Cx. pipiens* an ideal model for probing miRNA regulation of adult-reproductive diapause.

In this study we used quantitative reverse-transcript PCR (qRT-PCR) to measure candidate miRNAs that were selected because they have experimentally verified roles in diapause-relevant processes (e.g. circadian clock, ovarian development, insulin signaling, or metabolism) or are known to be differentially regulated in diapausing pupae of the flesh fly, *S. bullata* [30]. We measured miRNA abundance in diapausing and nondiapausing adult females 0, 5, 12, and 22 days after adult emergence, providing information about changes in miRNA profiles over the adult lifespan that may be related to reproductive development and aging. We also measured two phenotypic markers of diapause, egg follicle length and fat content, in female mosquitoes at these times. Finally, we measured changes in miRNA abundance in nondiapausing females after they were given a blood meal to determine the possible role of miRNAs in regulating processes related to blood-feeding and ovarian development in diapausing and nondiapausing *Cx. pipiens*. Together these data provide evidence that several miRNAs show diapause-related shifts in abundance and suggest inclusion of these small, noncoding RNAs in the diapause “toolkit”.

## 2. Methods

### 2.1 Insect rearing

The established laboratory colony of *Cx. pipiens* (Buckeye strain) was maintained as previously described [42, 48]. Mosquitoes in the main colony were kept at 25°C, 75% relative humidity under a long-day photoperiod (16 h light: 8 h darkness). Larvae were provided dried fish food (Tetramin; Blacksburg, VA USA). Adults were provided unlimited access to 10% sucrose solution. Chicken blood (Pel-freez Biologicals; Rogers, AR, USA) was provided using an artificial membrane system (Hemotek; Lancashire, UK) approximately 10 days post-adult emergence, and egg rafts were collected approximately 5 days later. To generate diapausing adults, larvae and pupae were held at 18°C, 75% relative humidity under a short-day photoperiod (8 h light: 16 h darkness). Diapausing adults had access to sugar water for the first 10 days of adult life and then sugar water was removed to simulate the lack of food in their natural environment. Nondiapausing adults used in these studies were generated by rearing all life stages under diapause-averting, long-day conditions (L:D 16:8) but at the same low temperature (18°C) with constant access to sugar water.

### 2.2 Measurement of ovarian development and total lipid content

The ovaries of 12 individual nondiapausing and diapausing females from each time point (0, 5, 12, and 22 days after adult emergence; n = 96 females total) were dissected in 0.9% saline solution, and the length of 10 egg follicles/female were measured under 200X magnification (Zeiss Axioskop, Thornwood, NY) as previously described [42].

The lipid content was measured in 4–5 diapausing and nondiapausing individual mosquitoes at the same developmental time points (0, 5, 12 and 22 days after adult emergence). In brief, dried mosquitoes were weighed, and the lipid content in each female was obtained using a modified Vanillin assay [52] as previously described [42] and divided by that female’s dry weight.

### 2.3 Sampling regime for miRNA expression studies

Changes in miRNA abundance were assessed in diapausing and nondiapausing females that had not been given a blood meal. Females were collected 0, 5, 12, and 22 d after adult emergence. All samples were collected 4 h after lights turned on (ZT4), and 4 biological replicates each containing 4–5 whole body, adult females were collected for each day.

Changes in miRNA abundance related to blood feeding, and possibly to stimulation of ovarian development, were measured in nondiapausing females 12 d after adult emergence. The experimental group was given a blood meal 36 h prior to sampling (i.e. were blood fed 10 d post-emergence). The control group, also sampled 12 d post-emergence, was only given sugar water and had never been given a blood meal. For each time point, there were 4 replicate samples of 4–5 females each. At the time of sampling, female mosquitoes were flash frozen and stored at −80 °C until they were processed.

### 2.4 Quantitative Reverse-transcription PCR

Abundance of candidate miRNAs was measured using qRT-PCR as previously described [30]. Total RNA was isolated from whole-body, female mosquitoes using the mirVana™ miRNA Isolation kit (ThermoFisher; Waltham, MA, USA) according to the manufacturer’s directions. Reverse transcription of 2 μg of total RNA was carried out using the miScript II RT kit (Qiagen; Valencia, CA, USA) according to the manufacturer’s directions for HiSpec buffer, which is designed to specifically transcribe mature miRNAs. Relative abundance of each candidate miRNA was measured using an iQ5™ Multicolor Real-time PCR Detection System (Bio-Rad; Hercules, CA, USA) and miScript Primer Assays (Qiagen) which use one universal primer and one primer designed to detect a specific miRNA sequence. Primer performance conformed to MIQE standards for efficiency [49] as shown in Supplementary Table S1. Cycling parameters were 94°C for 15 min followed by 40 cycles of 94°C for 15 s, 55°C for 30 s and 72°C for 30 s. Melt curve analysis indicated only one product was formed under these conditions.

Relative miRNA abundance was evaluated in 3–4 replicate samples for each group with three technical replicates for each miRNA assay using a modified 2^-ΔCt^ method [50]. Briefly, background subtracted fluorescence data were exported from the Bio-Rad iQ5 software and were smoothed and normalized as previously described [51]. These normalized data were used to determine the threshold cycle (Ct) for every technical replicate for each sample, and the average technical replicate for each biological sample was calculated. Extensive analysis demonstrated that the abundance of the miRNA let-7 was consistent across all of our samples (Supplementary Figure S1). Hence, the Ct values of the biological replicates were averaged and normalized by subtracting the average Ct for let-7 which served as an internal reference. The resulting value was log transformed to give the relative miRNA abundance (2^-ΔCt^).

### 2.5 Statistical analysis

All statistical analyses were performed in R.3.32 (R Core Team, 2017). Changes in average egg follicle length, lipid content, and miRNA abundance over time (i.e. day 0 to day 22) for a single mosquito type (i.e. diapausing or nondiapausing) were evaluated using a One-way ANOVA followed by Tukey’s post-hoc test. Differences between diapausing and nondiapausing females for each time point tested were evaluated using Student’s t-test, as were differences between sugar fed and blood fed females. To minimize type 1 errors, p-values for Student’s t-test were corrected using the False Discovery Rate (FDR) method [53].

## 3. Results

### 3.1 Changes in ovarian length and lipid content related to aging and diapause development

Arrested ovarian development and lipid sequestration are hallmark features of diapause in *Cx. pipiens* [33, 54]. Average egg follicle length (Fig. 1A), a measure of ovarian development, was modestly, but significantly, higher in females programmed to enter diapause than in nondiapausing females on the day of adult emergence (mean ± s.e.m = 47.4 ± 1.5 μm for diapausing females and 38.4 ± 6.1 μm for nondiapausing females; FDR-adjusted p < 0.001). In nondiapausing females, follicle length increased significantly during the next 22 days (One way ANOVA, p < 0.001). The largest increase occurred during the first 5 days with a significant 2.5-fold increase in average length (Tukey’s post hoc comparison Day 0 to Day 5, p < 0.001), followed by a smaller but significant increase in egg follicle length from days 12 to 22 (~1.2-fold; Tukey’s post hoc comparison Day 12 to Day 22, p < 0.001). There was a slight but significant increase in the egg follicle lengths in diapausing females over the first 22 days of adult life (~1.1 fold increase between 0 and 22 d post adult emergence; One way ANOVA, p <0.001). However, nondiapausing females had significantly larger egg follicles than diapausing females on days 5, 12 and 22 egg (FDR-adjusted p < 0.001 for each time point).

**Figure 1.**
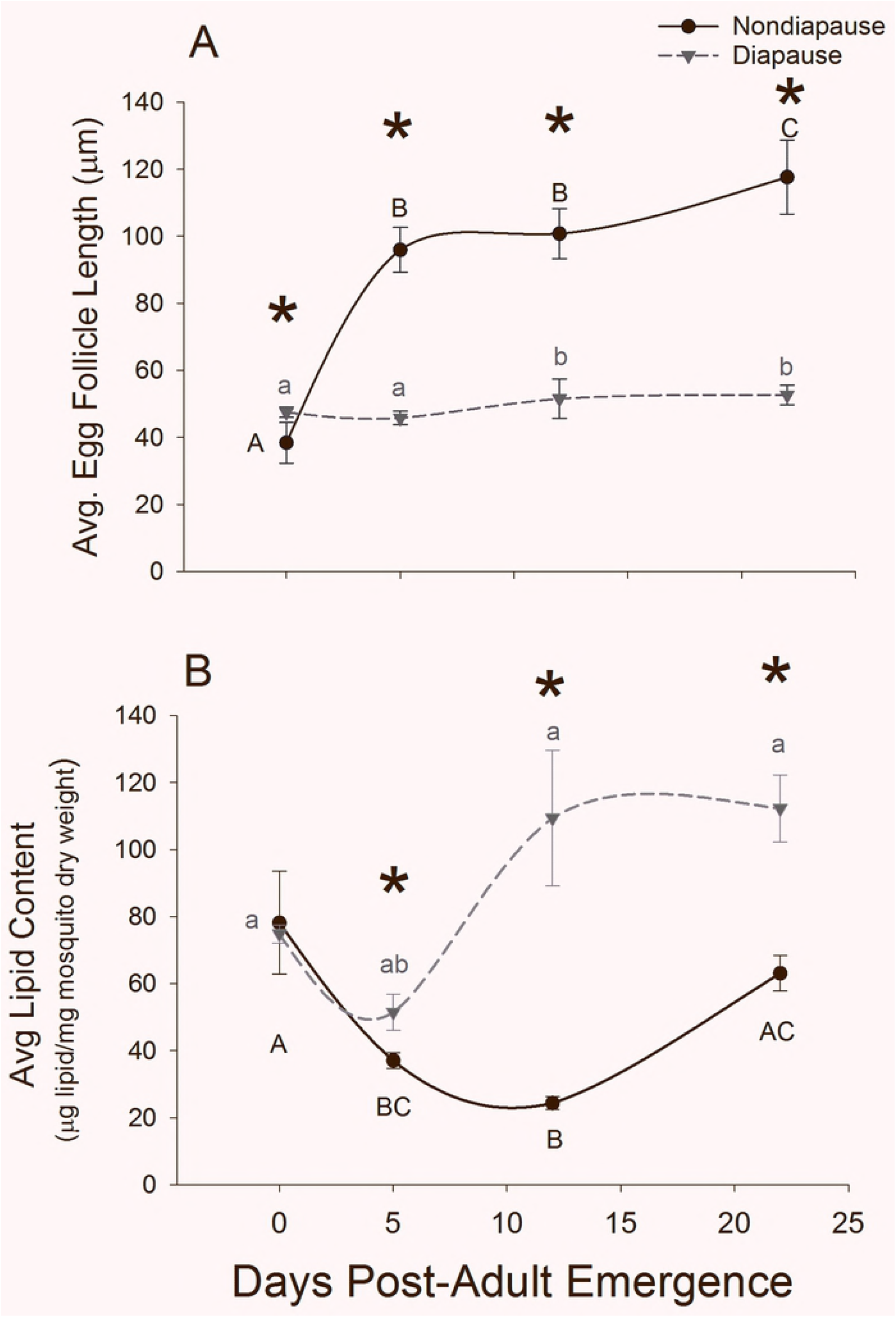
Changes in physical characteristics of diapausing and nondiapausing female mosquitoes. (A)Average egg follicle length of 12 females and (B) average lipid content in 4-5 females per time point and diapause status. Error bars represent s.e.m, and spline curves were fit to the data using SigmaPlot. Asterisks indicate significant differences in phenotypic markers between diapausing and nondiapausing females on the same day (Student’s T-test with Benjamini and Hochberg adjusted FDR, p < 0.05), while significant changes across the adult lifespan are represented with letters (black capital letters = nondiapausing mosquitoes; gray lowercase letters = diapausing mosquitoes; One-way ANOVA followed by Tukey’s Honest Significant Difference Test, p < 0.05).

There was no difference in average lipid content in pre-diapause females compared to females not programmed for diapause on the day of adult emergence (Fig. 1B; FDR-adjusted p = 0.840). In nondiapausing females, there was a significant decrease in total lipid/mg dry weight over the next 22 d (One way ANOVA, p = 0.001) with total reduction of approximately 60% between 0 and 12 d (Tukey’s post-hoc comparison, p = 0.001). In diapausing females total lipid content increased ~1.5 fold during the 22 d following adult emergence. Diapausing females had significantly more total lipid than their nondiapausing counterparts every day except the day of emergence (Day 0, 5, 12 and 22 FDR-adjusted p = 0.839, 0.0179, 0.048 and 0.0179 respectively).

### 3.2 Changes in miRNA abundance related to aging and diapause progression

This set of experiments evaluated abundance of candidate miRNAs in diapause and nondiapausing adult females on the day of adult emergence (0 d) and on several additional days as adults aged (days 5, 12, and 22). MiRNAs evaluated include miR-8-3p, miR-13b-3p, miR-14-3p, miR-124-3p, miR-275-3p, miR-277-3p, miR-289-5p, miR-305-5p, miR-309-5p, and miR-375-3p. The abundance of all of these miRNAs dynamically and significantly changed during the 22 d evaluated. In general, miRNA abundance was higher on day 0 and lower on day 22 indicating a general decrease in abundance as adults aged (Fig. 2).

**Figure. 2.**
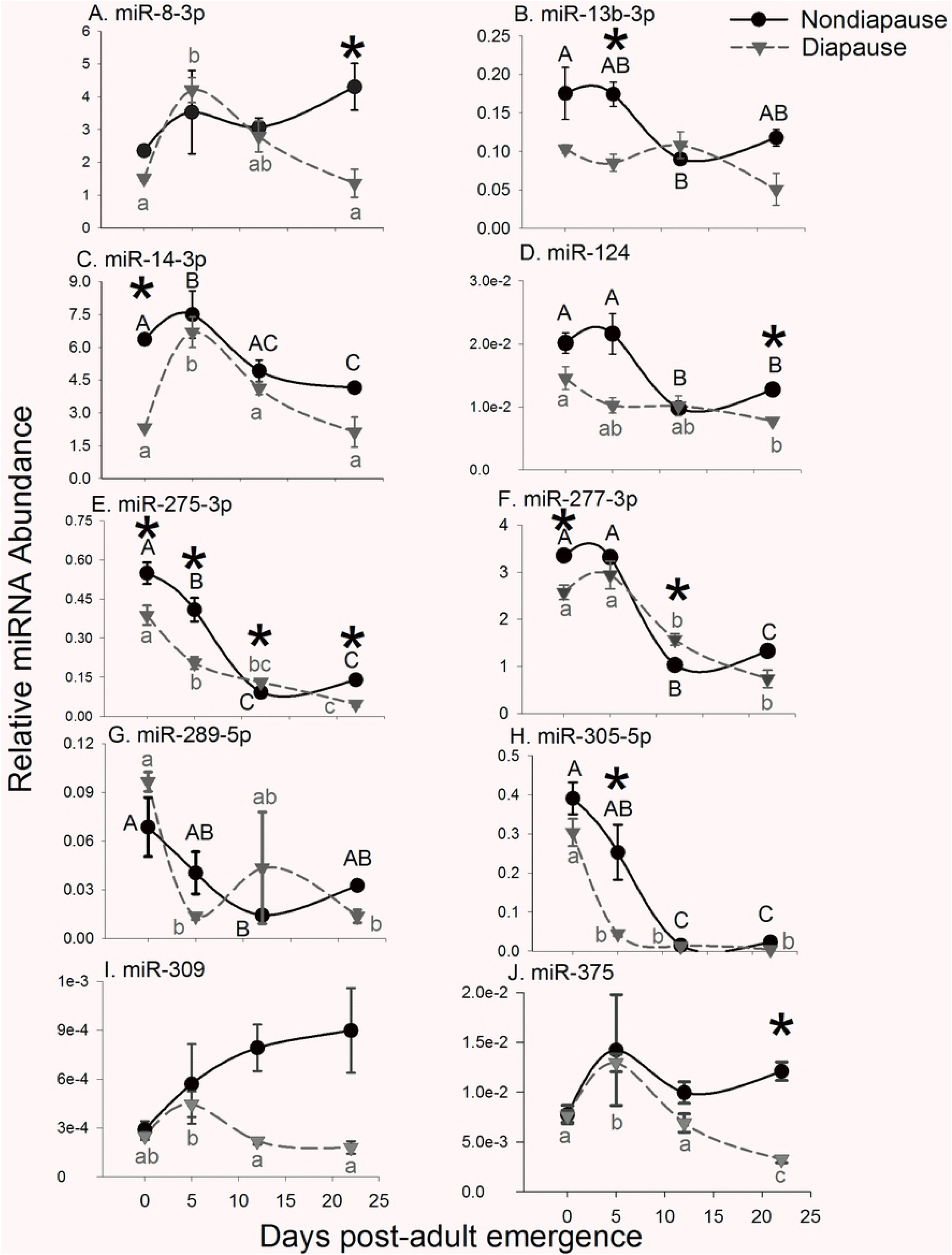
miRNA abundance dramatically changes in females across the adult lifespan and is frequently suppressed in diapausing mosquitoes. Relative abundance of (A) miR-8-3p, (B) miR-13-3p, (C) miR-14-3p, (D) miR-124-3p, (E) miR275-3p, (F) miR-277-3p, (G) miR-289-5p, (H) miR-305-5p, (I) miR-309-5p and (J) miR-375-3p measured with qRT-PCR. Each point represents the average relative miRNA abundance in 3–4 biological replicates each containing 3–5 whole female bodies. Bars represent s.e.m. Data were normalized to the microRNA let-7, and spline curves were fit to the data using SigmaPlot. Asterisks indicate significant differences in miRNA abundance between diapausing and nondiapausing females (Student’s T-test with Benjamini and Hochberg adjusted FDR, p < 0.05), while different letters indicate significant changes in miRNA expression across adult lifespan (black capital letters = nondiapausing females; gray lowercase letters = diapausing females; one way ANOVA followed by Tukey’s Honest Significant Difference test, p < 0.05;).

In nondiapausing females there was an apparent ~ 2-fold increase in miR-8-3p (Fig. 2A) abundance between 0 and 22 d post emergence that was not significant (One way ANOVA, p = 0.218). In diapausing females miR-8-3p abundance increased nearly 3-fold between 0 and 5 d followed by a decrease between days 5 and 22 (One Way ANOVA, p < 0.001; Tukey’s post-hoc comparison day 5 and day 22, p < 0.001). MiR-8-3p abundance was similar in diapausing and nondiapausing individuals except on days 0 and 22 when it was 1.5 and 3-fold more abundant in nondiapausing females than in diapausing females (Day 0 and 22 FDR-adjusted p = 0.0076 and 0.034, respectively).

There was a general decrease in the abundance of miR-13b-3p (Fig. 2B) in nondiapausing mosquitoes (One way ANOVA, p = 0.029) but no significant change in diapausing mosquitoes over 22 d (One way ANOVA, p = 0.069). MiR-13b-3p was approximately 1.8-fold more abundant in nondiapausing mosquitoes than diapausing mosquitoes on days 0 and 5 (FDR-adjusted p = 0.16 and 0.043, respectively) but there was no significant difference on days 12 or 22 (FDR-adjusted p = 0.38 and 0.080 respectively).

MiR-14-3p (Fig. 2C) significantly decreased ~1.5-fold in nondiapausing females (One-way ANOVA, p = 0.004). In diapausing females, there was a 3-fold increase between days 0 and 5 followed by a sharp decrease (One-way ANOVA, p < 0.001). On day 0 miR-14-3p was 4-fold more abundant in nondiapausing females compared to diapausing females (FDR-adjusted p < 0.001), but there were no significant differences between diapausing and nondiapausing females at any other time point.

MiR-124-3p (Fig. 2D) abundance significantly decreased from 0 to 22 d in both nondiapausing (One way ANOVA, p < 0.001) and diapausing (One-way ANOVA, p = 0.028) females. It was significantly more abundant in nondiapausing females only on day 22 (FDR-adjusted p <0.001). Although miR-124
appeared to be upregulated in nondiapausing females on days 0 and 5 (1.38 and 2-fold, respectively), these differences were not significant (FDR-adjusted p = 0.08 and 0.08 respectively).

MiR-275-3p (Fig. 2E) decreased significantly in both nondiapausing (One-way ANOVA, p < 0.001) and diapausing females (One-way ANOVA, p < 0.001). MiR-275 was moderately, but significantly, more abundant in nondiapausing females on days 0, 5 and 22 (FDR-adjusted p = 0.028, 0.028 and 0.002, respectively). MiR-275-3p was also significantly more abundant in diapausing females on day 12 (FDR-adjusted p = 0.01), but whether this difference is biologically meaningful is unclear.

Similarly, the abundance of miR-277-3p, miR-289-5p and miR-305-5p significantly decreased as females aged. Specifically, miR-277-3p (Fig 2F) decreased 2.5 fold in nondiapausing females (One way ANOVA, p < 0.001) and 3.5 fold in diapausing females (One way ANOVA, p <0.001) from day 0 to 22. MiR-277-3p was underexpressed in diapausing females on day 0 (FDR-adjusted p = 0.027) but was not significantly different from nondiapausing females at other times. MiR-289-5p (Fig. 2G) abundance significantly decreased over time in nondiapausing (One way ANOVA, p = 0.028) and diapausing females (One Way ANOVA, p = 0.018) but there were no significant differences in miR-289-5p abundance between diapausing and nondiapausing females. MiR-305-5p (Fig. 2H) decreased 17 fold in nondiapausing females (One way ANOVA, p < 0.001) and 80 fold in diapausing females (One way ANOVA, p < 0.001) from day 0 to day 22. MiR-305-5p was underexpressed in diapausing females on day 5 post-emergence (FDR-adjusted p = 0.013), but not at other time points.

MiR-309-5p abundance (Fig. 2I) did not significantly change in nondiapausing females during the 22 d following adult emergence (One way ANOVA, p = 0.147), but increased significantly in diapausing females on day 5 followed by a decrease to its initial level on days 12 and 22 (One way ANOVA, p = 0.008). Beginning on day 12, there was an apparent 3-fold difference between diapausing and nondiapausing females that was due to a substantial increase in miR-309-5p abundance that was not observed in diapausing females. However, this was not significant (FDR-adjusted p-values = 0.104 and 0.137 on days 12 and 22 respectively).

MiR-375-3p (Fig. 2J) abundance did not change significantly between 0 and 22 d following adult emergence in nondiapausing females (One way ANOVA, p = 0.306), but in diapausing females there was a significant increase in miR-375-3p between 0 and 5 d followed by significant decrease on days 12 and 22 (One Way ANOVA, p < 0.001). MiR-375-3p was ~3.7-fold more abundant in nondiapausing females than in diapausing females 22 d post-adult emergence (FDR-adjusted p = 0.004).

### 3.2 Target identification of diapause relevant miRNAs

The functional significance of differentially regulated miRNAs depends on the identity of the genes they regulate. Identifying gene targets of an individual miRNA is complicated because a single miRNA can regulate multiple mRNAs and a single mRNA can be regulated by multiple RNAs. To maximize our ability to identify diapause relevant targets of miRNAs, we used TargetScan Fly, release 6.2 [55, 56], to identify miRNAs that may regulate transcripts of candidate genes that are known, from previously published studies, to be differentially regulated in diapausing *Cx. pipiens* [8, 40]. We also used DIANA mirPath 3.0 [57] to identify KEGG pathways that may be regulated by miRNAs that were differentially regulated 0 and 5 d post-adult emergence.

Multiple studies have identified numerous genes that are differentially regulated in diapausing females of *Cx. pipiens*, many of which are putative targets of miRNAs evaluated in this study. Fifteen of the thirty-three genes that regulate fat metabolism in diapausing females of *Cx. pipiens* are predicted targets of miRNAs [38] (Supplementary Table S2). MiR-277-3p potentially regulates eleven genes including *acc* and *fabp*, which encode Acetyl-CoA carboxylase and Fatty acid binding protein, respectively. In addition, *fad-2* and *fad-3*, two genes which both encode Δ(9)-desaturase enzymes, are putative targets of miR-305-5p (*fad-2*) or miR-8-3p and miR-124-3p (*fad-3*)

A meta-analysis of diapause-relevant genes in insects [58] identified 572 *Drosophila* transcripts that are orthologous to diapause-relevant genes in *Cx. pipiens* [8]. Of these, 138 are putative targets of the miRNAs evaluated in this study. These include 39 putative targets of miR-277-3p, 21 targets of miR-375-3p, and 15 targets of miR-13b-3p (Table S2). Most of these putative targets were not differentially regulated in diapausing *Cx. pipiens* [8, 58]. However, miRNAs can repress translation without degrading the transcript [59], and it is possible for a gene to be regulated by a miRNA without a significant change in transcript abundance.

DIANA mirPath 3.0 was used to identify KEGG pathways that have genes that are regulated by specific miRNAs. We identified 8 KEGG pathways, including Mucin type O-Glycan biosynthesis; Valine, leucine and isoleucine degradation; Fatty acid elongation; Fatty acid degradation; Propanoate metabolism; MAPK signaling pathway; Hippo signaling pathway; and Valine, leucine and isoleucine biosynthesis, that may be regulated collectively by miR-14-3p, miR-275-3p, miR-13b-3p, miR-124-3p, miR-277-3p, miR-305-5p. It is important to note that all of these microRNAs were significantly lower in diapausing females 0 and/or 5 d post-adult emergence (Table 1), suggesting that these pathways are regulated as females prepare to enter diapause.

**Table 1.**
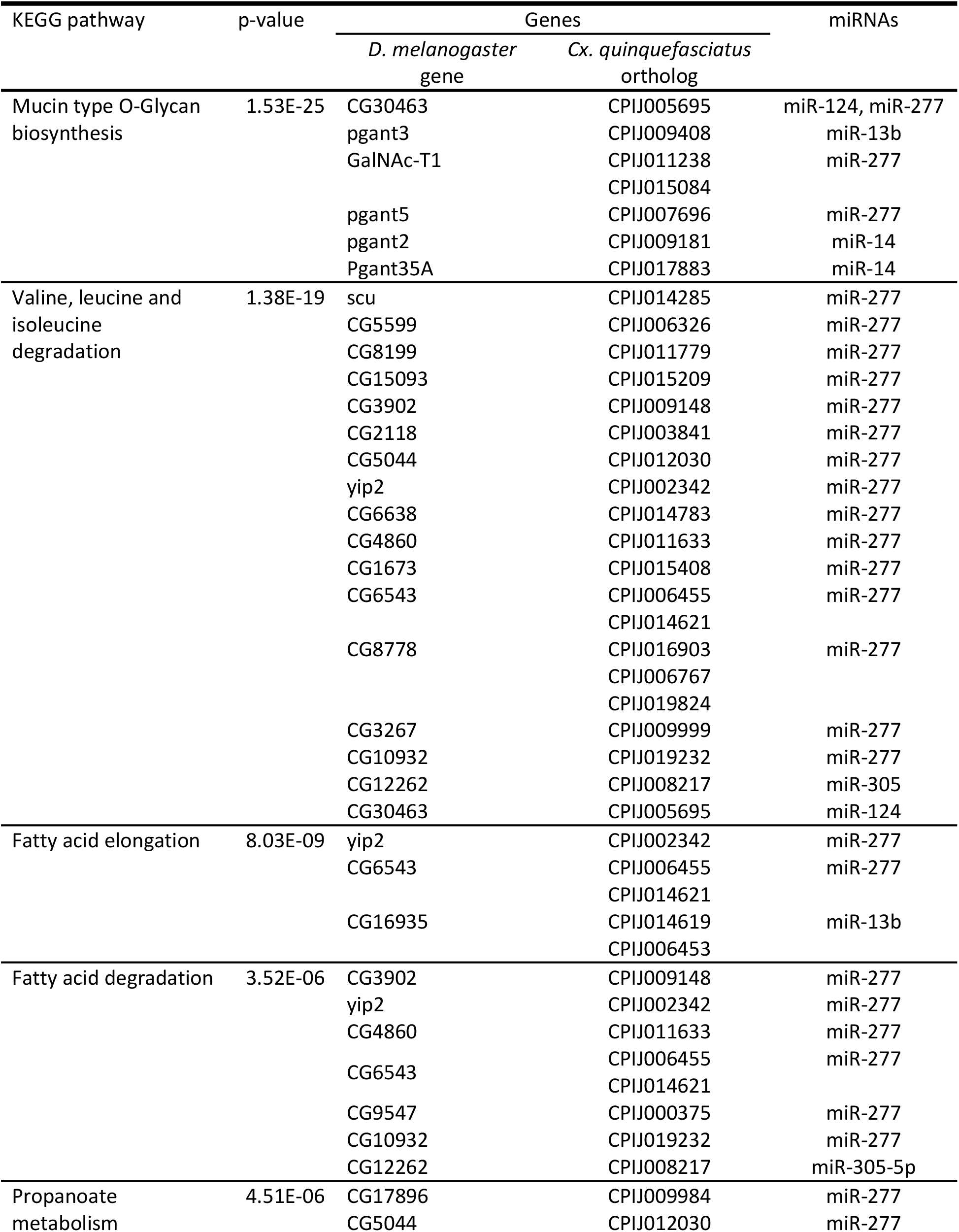
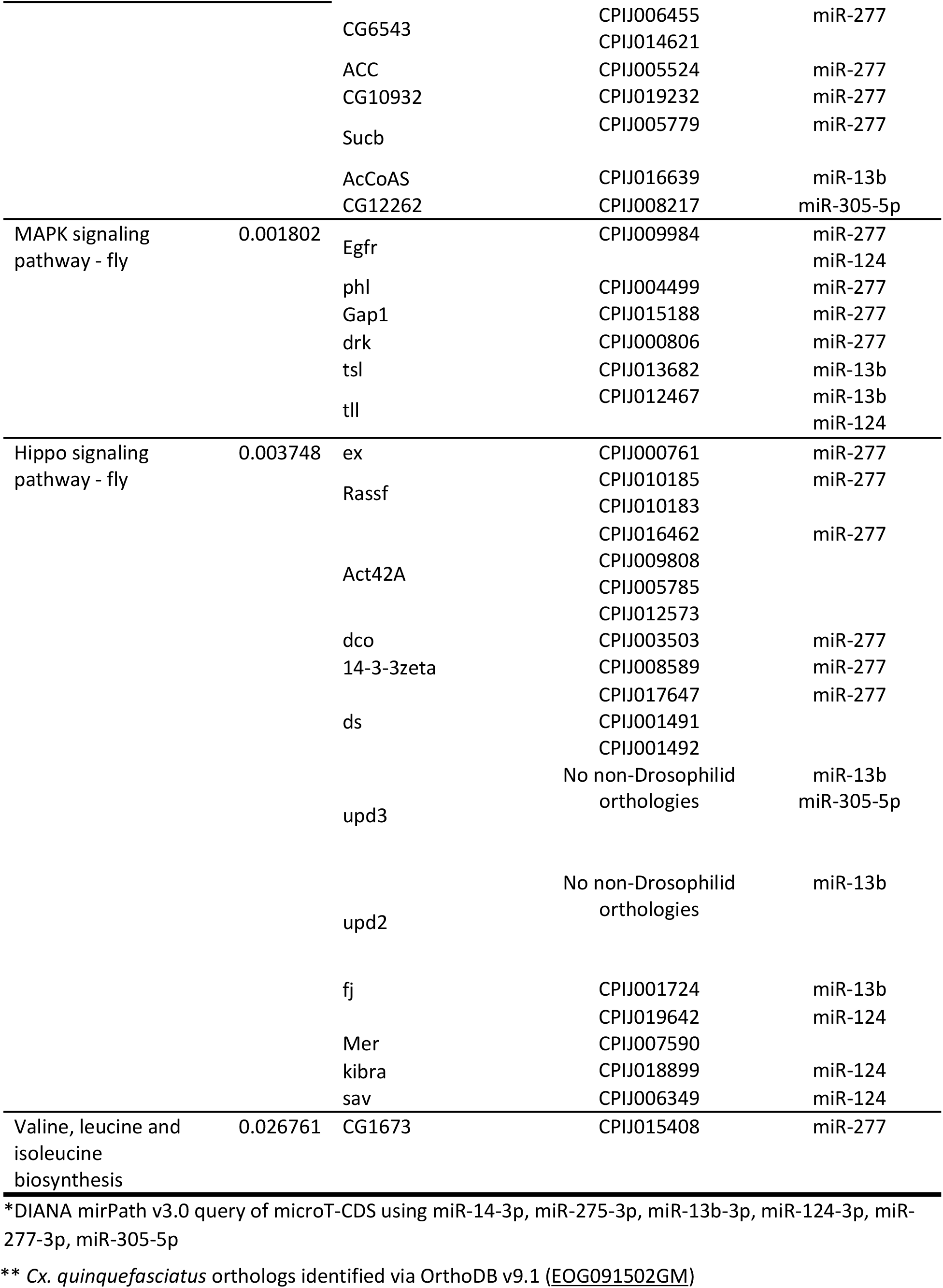
KEGG pathways that may include genes regulated by miRNAs that are differentially regulated before diapause entry*

### 3.4 Changes in miRNA abundance related to blood feeding

This set of experiments evaluated changes in abundance of candidate miRNAs known to be differentially regulated in female mosquitoes following a blood meal, which is required for egg production in most species (reviewed in [60]). MiR-124-3p, miR275-3p, miR-309-5p, and miR-375-3p were measured in long-day reared, nondiapausing females 12 days after adult emergence that had never received a blood meal (sugar-fed control) or had been given a blood meal 36 h prior to sampling. There was no significant change in miR-124-3p abundance (FDR-adjusted p = 0.102) but miR-275-3p, miR-309-5p and miR-375-3p were all significantly upregulated in blood-fed females (FDR-adjusted p = 0.021 for each).

**Figure 3.**
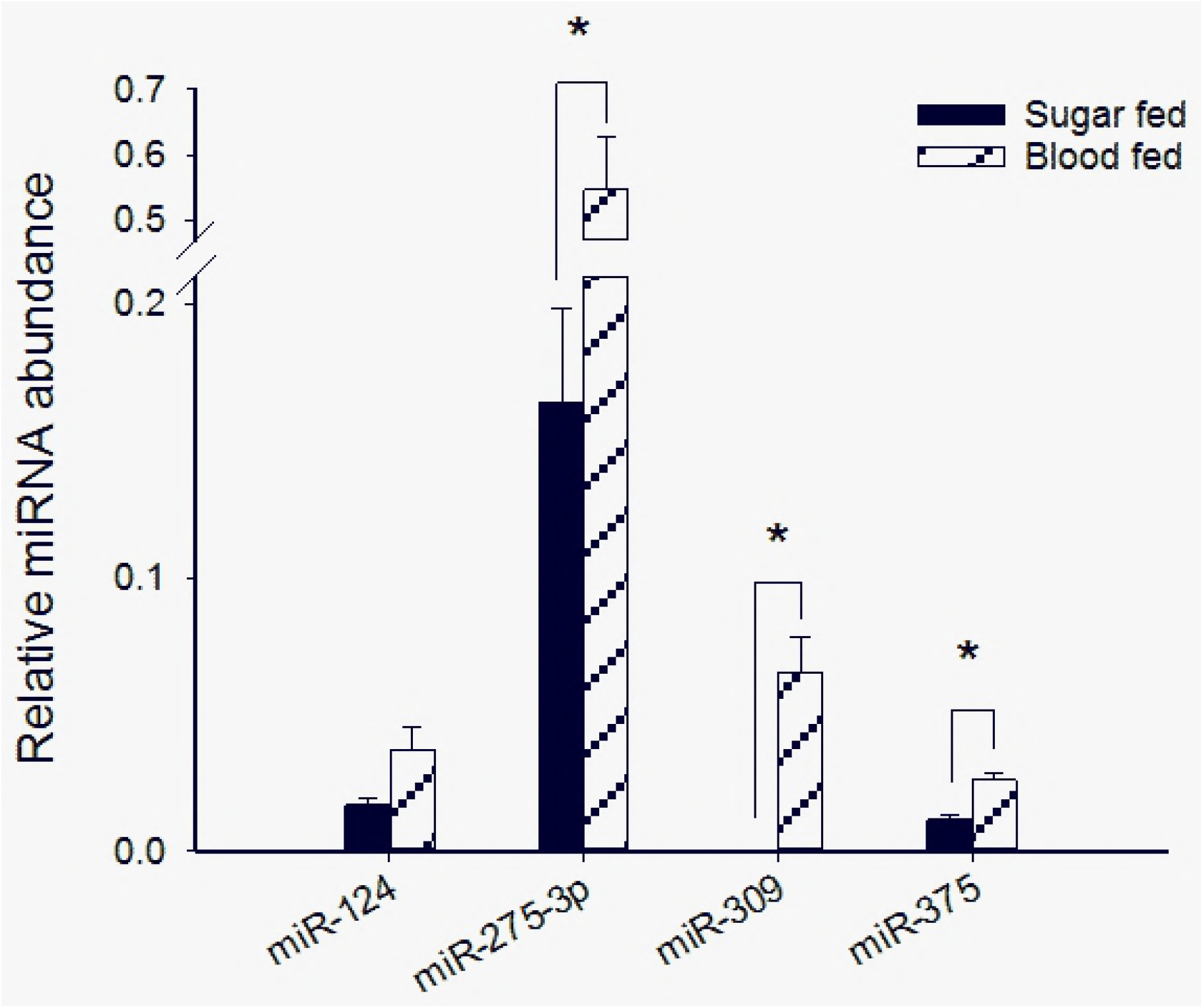
Several miRNAs are upregulated following a blood meal. Relative miRNA abundance was measured with qRT-PCR. Bars represent the mean ± s.e.m of in 4 independent, biological replicates each containing 3–5 whole body, nondiapausing females. All females were 12 days old and blood fed females ingested a blood meal 36 h before collection. Blood feeding did not significantly change the abundance of miR-124-3p (FDR-adjusted p = 0.102) but did significantly increase the abundance of miR-275-3p, miR-309-5p and miR-375-3p (FDR-adjusted p < 0.05).

## 4. Discussion

Diapause is a complex phenotype that depends on coordinated regulation of multi-gene networks that are themselves regulated by numerous factors, including microRNAs. The results of this study indicate that multiple microRNAs are differentially regulated in pre-diapause and diapausing females of *Cx. pipiens* mosquitoes compared to their nondiapausing counterparts. Specifically, miR-8-3p, miR-13b-3p, miR-14-3p, miR-275-3p, and miR-305-5p were underexpressed in diapause-destined females on days 0 and 5 following adult emergence and likely regulate diapause entry and/or the manifestation of phenotypes associated with diapause. MiR-8-3p, miR-124, miR-275-3p and miR-375-3p were underexpressed in diapausing females 22 d post-emergence and may have a role in diapause maintenance. Many of these miRNAs are also differentially regulated in diapausing pupae of *S. bullata* [30] indicating that microRNAs could be considered part of a conserved “toolkit” of mechanisms that mediate diapause entry and diapause maintenance in insects.

The functional significance of these changes in miRNA abundance, and how they regulate diapause, remains to be tested, but some inferences can be made based on computationally predicted gene targets and from studies on other insects including *D. melanogaster* and mosquitoes including *Aedes aegypti* and *Anopheles stephensi*. To facilitate the discussion, we focus on miRNAs that have known roles in circadian timing, lipid metabolism, and ovarian development, which are key processes in the establishment and maintenance of diapause in *Cx. pipiens* [42, 38, 33].

Diapause in *Cx. pipiens* is associated with changes in the expression profiles of core components of the circadian clock [42]; and miR-124-3p, expressed primarily in the brains of mosquitoes [61], may regulate some of these changes. In *D. melanogaster*, miR-124-3p regulates circadian output by targeting the gene encoding the CLOCK protein [62–64]. MiR-124-3p also mediates daily phases of locomotor activity by regulating genes in the Bone Morphogenetic Protein signaling pathway [65, 66]. Therefore, we think that miR-124-3p likely regulates behavioral and physiological outputs downstream of the circadian clock in diapausing *Cx. pipiens*. One possibility is that reduced miR-124 in diapausing mosquitoes drives changes in sugar feeding activity of mosquitoes that have previously been reported [67]. It is important to note, however, there are some fundamental differences in the regulation of the circadian clock in *Cx. pipiens* compared to *D. melanogaster* [68], and additional studies are needed to confirm the link between miRNAs, circadian rhythms, and diapause in *Cx. pipiens*.

Diapause in *Cx. pipiens* is characterized by fat accumulation, through overall suppression of insulin signaling [40, 69]. We found that, on the day of adult emergence (i.e., day 0) there was no difference in the total lipid content of short-day reared, diapause-destined females compared to nondiapause-destined females. However, there was a significant, 1.5 fold (FDR-adjusted p = 0.006) difference between the two groups by day 5, indicating rapid fat accumulation occurs during this time. Indeed, this is consistent with previously reported changes lipid accumulation in diapausing females [54] and is also consistent with changes in mRNA expression for several genes associated with fat metabolism 1 week after adult eclosion [38]. Three miRNAs, miR-14-3p, miR-277-3p, and miR-305-5p that are known to regulate fat metabolism and/or insulin signaling [70–73], were downregulated in diapause-destined mosquitoes during this time, and, thus, are thought to contribute to the observed changes in lipid metabolism in mosquitoes entering diapause.

MiR-277-3p regulates insulin signaling directly by targeting genes that encode Insulin-like peptides (Ilps) [74] and indirectly by regulating branched-chain amino acid (i.e. valine, leucine, and isoleucine) metabolism [71]. In *Ae. aegypti* mosquitoes, knockdown of miR-277-3p reduces lipid storage in the fat body by upregulating insulin signaling and promoting nuclear export of the FOXO transcription factor [74–75]. In *D. melanogaster*, miR-277-3p indirectly mediates insulin signaling through regulation of branched-chain amino acid (BCAA) metabolism [71]. BCAAs activate TOR signaling and insulin secretion and regulate lifespan in evolutionarily diverse species [71, 76, 77]. Indeed, valine, leucine and isoleucine biosynthesis was one of the KEGG pathways that our computational analyses predicted to be regulated by miR-277-3p and other microRNAs that we observed were downregulated early in diapause. Taken together, these studies in *Ae. aegypti* and *D. melanogaster* and our results suggest that miR-277-3p regulates multiple aspects of insulin signaling and fat metabolism during diapause in *Cx. pipiens* and may be a critical player in the generation of the fat hypertrophy that is a hallmark of diapause in this species.

MiR-305-5p also regulates insulin signaling and lifespan in *D. melanogaster* [72, 73] and may mediate lipid accumulation during diapause in *Cx. pipiens* along with miR-277-3p. Six genes that are computationally predicted targets of miR-277-3p and miR-305-5p belong to the Fatty acid elongation and Fatty acid degradation pathways and are differentially regulated in diapausing females of *Cx. pipiens* 7 days post-emergence [38]. One gene of particular interest is *acd-1*, a putative target of miR-305-5p, which encodes an Acyl-CoA dehydrogenase and is overexpressed by ~2-fold in diapausing females 7 d post-emergence [38]. Suppression of miR-305-5p on day 5 is consistent with upregulation of *acd-1* two days later. However, the putative interaction of miR-305-5p and *acd-1* is based on the sequence of this gene in *D. melanogaster*, and additional work is needed to confirm that miR-305-5p regulates *acd-1* in *Cx. pipiens*.

In *D. melanogaster*, miR-14-3p suppresses insulin signaling by inhibiting its target *sugarbabe*, such that low levels of miR-14 lead to an increase in the level of insulin-like peptides [78]. Mutant flies lacking miR-14-3p store fat while flies overexpressing miR-14-3p are lean [70, 78]. Lower levels of miR-14-3p on day 0 in diapause-destined flies could indicate that miR-14-3p also regulates fat metabolism in *Cx. pipiens*. However, it is difficult to rationalize the changes in miR-14-3p abundance in diapausing and nondiapausing females, as insulin signaling is suppressed in diapausing females [40] while lipid content is elevated (Fig. 1). In addition, it is not clear whether the function of miR-14-3p in *D. melanogaster* is conserved in other insects. The 3′ UTR sequence of *sugarbabe* that is available for *Cx. quinquefasciatus* (CPIJ007837) does not appear to have a region that miR-14-3p can bind. However, there are multiple regions within the open reading frame (ORF) of *sugarbabe* where miR-14-3p could potentially bind. It will be interesting to see whether the interaction between miR-14-3p that occurs in *D. melanogaster* is conserved in mosquitoes and whether miR-14-3p regulates insulin signaling and fat metabolism in diapausing females of *Cx. pipiens*.

Arrested ovarian development is also a key feature of the diapause phenotype in *Cx. pipiens* [33]. Surprisingly, diapause-destined females had significantly larger egg follicles than nondiapausing females on the day of adult emergence (i.e. day 0). A significant increase in egg follicle length in nondiapausing females, but not in diapausing females, between days 0 and 5 suggests that ovarian development is suppressed in diapausing females during this time. Numerous miRNAs have experimentally validated roles in ovarian development in mosquitoes, including miR-8-3p, miR-275-3p, miR-309-3p, and miR-375-3p [43, 45, 79–80]. Of these, only miR-8-3p and miR-275-3p were differentially expressed early in diapause entry. MiR-8-3p, miR-275-3p, and miR-375-3p were all significantly underexpressed in diapausing females on day 22 post-emergence, which suggests these miRNAs, or their absence, is important for continuously arresting egg follicle development and/or other pathways that maintain diapause in *Cx. pipiens*.

It is important to note that these miRNAs were differentially regulated in diapausing and nondiapausing females that had only been given sugar water and had never taken a blood meal. Reports from other mosquito species that also require a blood meal for ovarian maturation showed that expression of these miRNAs increases substantially following a blood meal [43, 45, 79–81] and, indeed, we found the abundance of miR-275-3p, miR-309-3p, and miR-375-3p increased by as much as 193-fold after females of *Cx. pipiens* consumed blood. However, there are significant differences in the baseline levels of these miRNAs, and in egg follicle length, between diapausing and nondiapausing females in the absence of a blood meal (Fig. 1A). Therefore these miRNAs are likely suppressed as part of the diapause program, independent of blood feeding.

Of the miRNAs that regulate ovarian development, miR-8-3p is of particular interest because it also targets genes involved in fat metabolism. In *D. melanogaster*, miR-8-3p regulates insulin signaling by inhibiting translation of U-shaped, a protein that activates phosphatidylinositol 3-kinase (PI3K) [82]. The 3′ UTR is not available for the *Cx. pipiens* ortholog of U-shaped, so it is not known whether miR-8-3p regulates this gene in this mosquito species. In *Ae. aegypti* miR-8-3p is expressed in the fat body and coordinately regulates both fat metabolism and reproduction by targeting *secreted wingless-interacting molecule* (*swim*), a gene in the Wingless signaling pathway [79]. The *Culex* ortholog of *swim* (CPIJ008716) contains a region in the 3′ UTR where miR-8-3p can bind (Supplementary Fig. S2), thus it is likely this miR-8-3p function observed in *Ae. aegypti* is conserved in *Culex* mosquitoes. It will be interesting to further investigate how miR-8-3p regulates adult reproductive diapause in *Cx. pipiens*.

Taken together, our results demonstrate that miRNA expression in diapausing and nondiapausing females of *Cx. pipiens* mosquitoes is dynamic and changes as adults age. We observed that nearly all miRNAs examined were less abundant in diapausing females than in nondiapausing *Cx. pipiens* at some point in development. However, these differences varied throughout adulthood, which further underscores the dynamic nature of miRNA expression during reproductive development and diapause progression. Our results also demonstrate that diapause entry is associated with changes in miRNA abundance in females of *Cx. pipiens*. Although it is possible that the diapause phenotype is responsible for these differences, our *in silico* analyses have uncovered several diapause-relevant genes and pathways that are likely regulated by the microRNAs we have examined. Therefore, we provide compelling evidence that microRNAs are likely generate and maintain the diapause phenotype. The complex nature of miRNA gene regulation, where each miRNA has hundreds of potential mRNA targets that it can either positively or negatively regulate, leads to a dizzying number of possibilities in how the differences in miRNA abundance we have identified here might impact the diapause program. Future experimental assays that manipulate miRNA levels are essential to clarify the functional role of these small RNAs in the diapause “toolkit”. Such experiments will likely reveal how mosquitoes and other animals use miRNAs to translate environmental signals into physical molecular regulators to appropriately coordinate their seasonal growth, development, and reproduction.

## 5. Acknowledgements

We thank David Denlinger for experimental advice and reviewing an early draft of this manuscript. We also thank Drew Spacht and Justin Peyton.

## Supplementary Figure Captions

Supplementary Table S1. Sequence of each miRNA-specific primer and its efficiency (E) and regression coefficient (R^2^) from standard curve analyses.

Supplementary Table S2. Putative miRNA target genes involved in fat metabolism as identified by Sim and Denlinger, 2009.

Supplementary Table S3. Putative miRNA targets in diapausing *Cx. pipiens*. Adapted from data collected by Kang et al., 2015 and analyzed by Ragland and Keep, 2017.

Supplementary Figure S1. Consistent cycle threshold (C_t_) values for let-7 demonstrate that it is a viable reference gene. (A) Abundance of let-7 in diapausing and nondiapausing females collected on days 0, 5, 12 and 22 (n = 31); (B) abundance of let −7 sugar fed and blood fed nondiapausing female mosquitoes (n = 8).

Supplementary Figure S2. Interaction between *swim* and miR-8-3p. The seed region of miR-8-3p (shown in red font) aligns perfectly with the 3′ UTR of *swim* and binding between miR-8-3p and *swim* is energetically favorable. Taken together, this suggests that miR-8-3p likely inhibits translation of *swim* in *Culex* mosquitoes.

